# Non-dichotomous inference using bootstrapped evidence

**DOI:** 10.1101/017327

**Authors:** D. Samuel Schwarzkopf

**Affiliations:** Experimental Psychology, University College London, 26 Bedford Way, London WC1H 0AP, United Kingdom; Institute of Cognitive Neuroscience, University College London, 17 Queen Square, London, WC1N 3AR, United Kingdom

## Abstract

The problems with classical frequentist statistics have recently received much attention, yet the enthusiasm of researchers to adopt alternatives like Bayesian inference remains modest. Here I present the *bootstrapped evidence test*, an objective resampling procedure that takes the precision with which both the experimental and null hypothesis can be estimated into account. Simulations and reanalysis of actual experimental data demonstrate that this test minimizes false positives while maintaining sensitivity. It is equally applicable to a wide range of situations and thus minimizes problems arising from analytical flexibility. Critically, it does not dichotomize the results based on an arbitrary significance level but instead quantifies how well the data support either the alternative or the null hypothesis. It is thus particularly useful in situations with considerable uncertainty about the expected effect size. Because it is non-parametric, it is also robust to severe violations of assumptions made by classical statistics.

## Introduction

To this day classical null hypothesis significance testing remains the dominant approach for inferring the validity of an observed result in the psychological and life sciences. It rests on the probability (‘p-value’) that the observed effect, or a more extreme one, could have occurred under the assumption that there is no population effect. If the p-value is sufficiently low, this null hypothesis is rejected. However, p-values are frequently misinterpreted by researchers, uninformative about the evidence *for* an experimental hypothesis, highly susceptible to biased data sampling strategies, and generally prone to false positives [1–11]. The most devastating effect of p-values may be that they encourage an artificial dichotomy between significant and non-significant results[5].

The scatter plots in Fig. 1 illustrate the problems with p-values. Fig. 1A shows an almost perfect correlation between two measures (r=0.995, p<0.0004). However, there are only five observations. In contrast, the data in Fig. 1B are clearly correlated even though the correlation is weaker. Yet the p-value is similar (r=0.535, p<0.0004) because the sample size is much larger. Surely the evidence for a correlation in the second example is more compelling and more likely to replicate?

**Fig. 1.**
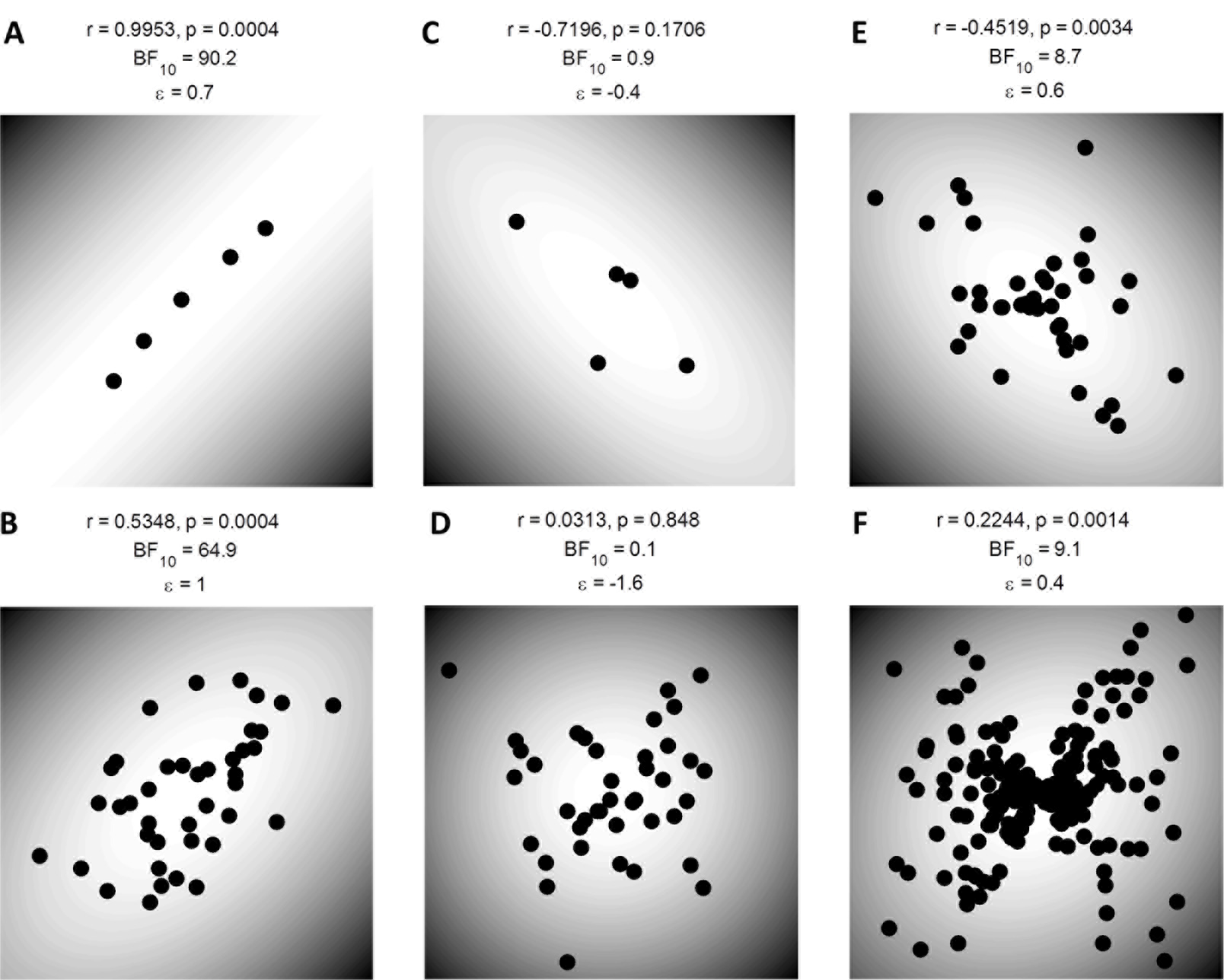
Scatter plots showing examples of correlation analysis. A-B. Correlated Gaussian data with n=5 (A) and n=40 (B). C-D. Uncorrelated Gaussian data with n=5 (C) and n=40 (D). E-F. Severely heteroscedastic data with n=40 (E) and n=200 (F). Each black dot is one observation. The grey shading denotes the Mahalanobis distance from the bivariate mean. Above each panel the Pearson’s correlation coefficient, the default Bayes factor [26] BF_10_ comparing the alternative and the null hypothesis, and the bootstrapped evidence ε are given.

One journal went so far as to ban the use of classical inference completely while proposing no viable alternative [8]. Others proposed guidelines to focus on effect size estimation and confidence intervals instead [5,12]. However, the use of confidence intervals is also fraught with problems [13] and may simply become a new significance testing procedure in disguise [5,14]. Moreover, like p-values, confidence intervals are frequently misinterpreted [14] and may perform inadequately [15]. Evidence for a hypothesis should *compare* an experimental (alternative) hypothesis to a baseline (null) hypothesis. Bayesian hypothesis tests using Bayes factors can achieve that but are often difficult to apply and rely on the choice of a prior, which can result in considerable debate (see e.g. [3,16–19]).

Here I present the *bootstrapped evidence* (BSE) test. It makes minimal assumptions and is applicable to a wide range of situations. Crucially, it quantifies the evidence for either the alternative or the null hypothesis non-dichotomously. Yet unlike Bayesian methods it is based only on the existing data without any question about prior distributions. It works by bootstrapping the effect size distributions under the two hypotheses by using different resampling strategies for each. For instance, when quantifying evidence for a linear correlation between two variables as in Fig. 1, under the null hypothesis data are resampled without respect to how individual observations have been paired. In contrast, under the alternative hypothesis the pairing is held intact but pairs are resampled to estimate the strength of the correlation. The evidence measure quantifies how distinct the distributions for the two hypotheses are and also incorporates the precision of the estimates. Thus, rather than determining probabilities as most inferential methods, it provides a *signal-to-noise ratio* for the hypothesized effect. Large evidence suggests a strong effect relative to the uncertainty with which it can be estimated.

Simulations demonstrate the test’s efficacy and robustness and compare it to classical frequentist and Bayesian inferential methods. Further, I apply this test to several concrete examples to show its advantages in practice. The MATLAB source code and example data are available for download (http://dx.doi.org/10.6084/m9.figshare.1342798) and we also plan to publish a standalone interface-based version and source code for R at a later date.

## Methods

I will refer to the distributions for the effect size under null and alternative hypothesis as θ_0_ and θ_1_, respectively (Fig. 2, left panels, blue and red curves). To quantify the *similarity* of these two distributions at each bootstrapping step the difference between the effects under the two hypotheses is calculated, that is, θ_1i_ - θ_0i_ where *i* denotes the *i*-th bootstrap step. This produces a *distribution of differences, Δ,* between the effects expected under the two hypotheses (Fig. 2, right panels). This is necessary because either θ_0_ or θ_1_ can be skewed separately by anomalous data. The Δ distribution captures either of these distortions.

**Fig. 2.**
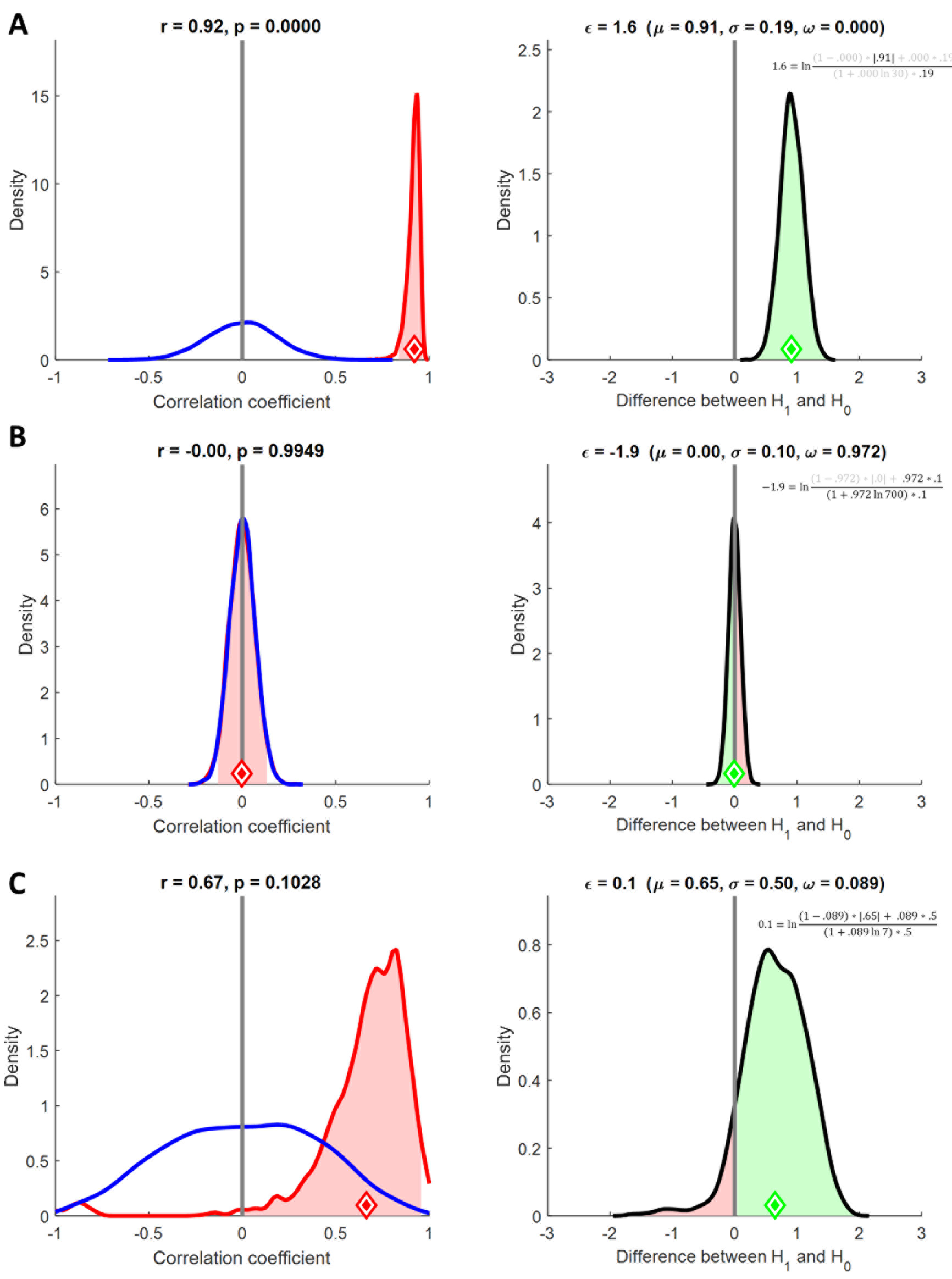
Examples of the bootstrapped evidence procedure. *Left panels* show the effect size distribution under the null (blue) and alternative (red) hypothesis estimated by bootstrapping. The red diamond denotes the observed effect size. The red shaded region denotes though the 95% confidence interval of the estimated effect. *Right panels* show the distribution of differences, Δ, between the alternative and null distributions (see Methods). The green diamond denotes the mean, μ. The ratio of the red and green areas under the curve is ω and quantifies the overlap between the null and alternative distributions. The standard deviation of Δ is denoted by σ. The bootstrapped evidence, ε, incorporates these three parameters and the sample size n and is expressed on a logarithmic scale (see equation). A. Strongly correlated data with n=30. Evidence compellingly supports H_1_. B. Uncorrelated data with n=700. Evidence compellingly supports H_0_. C. Uncorrelated data with n=7. Evidence is inconclusive.

Theoretically, the evidence for H_1_ is a standardized score based on the Δ distribution:

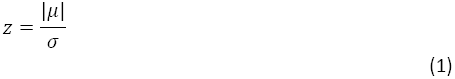

where μ and σ, respectively, denote the arithmetic mean and standard deviation of Δ. This effectively normalizes the shift of this distribution relative to zero by its dispersion. When z is greater than 1 this implies that the sample effect size, and thus our estimate of the true population effect, is larger than the *uncertainty* of the estimate (Fig. 2A). This provides evidence supporting the alternative hypothesis. The more data we collect and the sample size, n, becomes larger, the more accurate is the estimate of the population effect, μ, and the smaller is the uncertainty, σ.

Unfortunately, this only holds when the alternative hypothesis is true. When the null hypothesis is true instead, that is, when the population effect is zero, the parameters of Δ do not reflect the evidence for H_0_. While increasing n reduces the uncertainty, σ, on average it also reduces estimates of the population effect size, μ. It follows that the ratio of these parameters, z, remains more or less constant.

Thus, in order to quantify how strongly the data support *either* H_1_ or H_0_ we must weight the ratio of μ and σ according to how much evidence is available. This is achieved by calculating a third parameter describing the overlap of θ_0_ and θ_1_. This is given by:

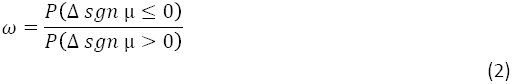

This divides the proportion of bootstraps in Δ that have the opposite sign as μ or zero (red area under the curves in the right panels of Fig. 2) by the proportion of iterations with the same sign (green area under the curves in the right panels of Fig. 2). The strength of evidence, ε, for or against H_1_ is then calculated as:

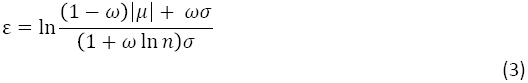

While this equation may seem complex, it is essentially the same as z, the standardized score in equation 1, but it is moderated by the sample size and the overlap between θ_1_ and θ_0_. When the data clearly support the alternative hypothesis, the Δ distribution is shifted far away from zero and thus ω (the ratio of the red and green areas under the curve) is very small (Fig. 2A). In fact, because it is based on the proportion of bootstraps it may equal 0. Under these circumstances the denominator is close to σ, the numerator is close to |μ|, and thus the evidence is approximately z from equation 1. However, when the null hypothesis is true, or the effect size is too small to clearly support the alternative, the Δ distribution is centered near zero and thus ω is near 1. In this situation the denominator is a multiple of σ, growing ever larger as the sample size increases. This in turn ensures that the ratio in equation 3 becomes ever smaller and evidence for the null hypothesis grows (Fig. 2B).

The numerator is also moderated by the strength of the evidence. When H_0_ is true and ω is near 1, the numerator is close to σ. This reflects the fact that when the data provide only weak evidence, there is substantial uncertainty as to whether the estimate of the effect size is accurate. When the sample size is large, this means that the denominator dominates the equation and ε becomes very small. However, when the sample size is *small*, the ratio in equation 3 is close to 1 and thus the data support neither H_1_ nor H_0_ very clearly – this means the evidence is inconclusive (Fig. 2C).

To recap, the bootstrapped evidence, ε, measures how confident one can be of the effect size estimate given the uncertainty in the data. If the observed effect is relatively large, the uncertainty will decrease as sample size grows and thus support for the alternative hypothesis also grows. However, when the effect remains considerably smaller than the uncertainty, a larger sample instead provides greater evidence for the null hypothesis.

Finally, as equation 3 shows, ε is the *natural logarithm* of this ratio. Therefore, when the ratio is near 1 and the evidence is inconclusive, ε is approximately 0. Positive ε indicates evidence for H_1_, while negative ε indicates evidence for H_0_. The bootstrapped evidence thus provides a non-dichotomous measure of the evidence for either hypothesis, similar to a Bayes factor [2,20,21]. However, while a Bayes factor is a measure of how much one should update the prior odds due to the observed evidence, the bootstrapped evidence is in essence a signal-to-noise ratio. A strong “signal” implies strong evidence for H_1_, while a negligible signal with a lot of data provides strong evidence for H_0_. The only prior assumptions this procedure makes pertain to how the data are resampled under the two hypotheses.

### Bootstrapped correlations

To bootstrap the evidence for a linear correlation data are resampled *with replacement* and on each step the correlation coefficient is computed. To derive the null distribution (θ_0_) data are resampled without restriction as would be expected if the effect occurred by chance, that is, observations for the two variables are no longer paired but intermixed randomly. This is essentially standard procedure for non-parametric resampling methods in the classical frequentist framework (although for this purpose permutation analysis where resampling is performed *without* replacement is more common). A classical one-tailed p-value could be calculated by determining the proportion of bootstraps θ_0i_ that are at least as large as the observed effect size – that is, the area under the blue curve to the right of the red diamond.

However, to derive the alternative distribution (θ_1_), quantifying the reliability of the observed effect, we restrict the resampling strategy on the alternative hypothesis that there is a correlation. In this case the pairing of data points in each variable is preserved so many resamples will show a positive linear relationship.

As described, we next calculate Δ, the distribution of differences between θ_1_ and θ_0_ (Fig. 2B). Its standard deviation is the uncertainty, σ. For all of the examples given in this article, I used 10,000 bootstrap iterations, except in the interest of time for lengthy simulations and curve fitting examples I only used 1,000 iterations. Reducing the number of iterations only changes the precision of the estimate of ε but does not alter the general conclusions substantially.

The further apart the two distributions for θ_0_ and θ_1_ are, the farther Δ is from zero and the greater is the evidence for H_1_. This is quantified by ε. When ε is very negative, the evidence favors H_0_ because it means the two distributions for θ_0_ and θ_1_ overlap considerably which means that Δ is centered near zero. Intuitively this indicates that the effect size estimate under H_1_ could very likely have been smaller than that under H_0_. When the evidence is -0.5<ε<0.5 this provides inconclusive support for the either hypothesis. This region is somewhat arbitrary but it reflects the fact that while there is overlap between the distributions for θ_0_ and θ_1_, there is not sufficient data to be confident that there is no subtle effect. This corresponds to the range of ε one typically obtains with small sample sizes when the null hypothesis is true.

### Bootstrapping differences

Naturally, the same procedure can be applied to other statistical comparisons in addition to tests of correlation. For instance, when *comparing the means of two independent samples* the effect size is the difference between the sample means. To estimate the null distribution, θ_0_, observations are resampled with replacement and divided into new samples of the same size as the original samples. To estimate the distribution for the alternative hypothesis, θ_1_, the segregation of the two samples is maintained and resampling is done *within* each sample. In all other respects, the procedure is identical to the correlation test already described.

When testing the *difference between two repeated measures* the same underlying principle applies. Here the effect size is the mean over the *differences* in each pair of observations. We estimate θ_1_ by maintaining the pairing but resampling with replacement. For estimating θ_0_ the pairing is also kept intact because what matters is only the variance across repeated measures. However, under the null hypothesis the order of measures is irrelevant so the resampling *randomizes the sign* for each observation. This corresponds to scrambling the order of observations in a repeated measures design.

Similarly, the BSE can also be used to test the *difference between two correlations*. Again the data need to be resampled based on the assumptions of the null as well as the alternative hypothesis. In this case the null hypothesis resamples data ignoring how variables are paired while the alternative hypothesis preserves the pairing. The estimated effect size is the difference between the two correlation coefficients.

The situation becomes more complicated for testing the *difference of one sample from a fixed value* (e.g., when comparing a normative sample to a patient case-study). Conceptually, it is possible to use the same resampling strategy as for a repeated measures design: the observations are the differences from the fixed value and for resampling the null distribution θ_0_ we randomize the sign of each observation. However, this approach is probably not sufficiently conservative. While it is conceptually correct for repeated measures designs to assume a mean difference of 0 under the null hypothesis, in many other situations fixed values are themselves subject to variability and/or measurement error. For instance, a measurement in a case study is subject to within-subject variability and chance performance in a behavioral task follows a probability distribution. Using a similar approach one can also incorporate variability in individual observations to calculate a group mean. In each round of bootstrapping we can simulate a new sample by drawing random data using the individual means and variances for every observation. This approach may be particularly suitable for meta-analysis.

### Bootstrapping tests against chance

Simulating the null distribution is also suitable for testing binomial processes, such as whether a coin is fair or whether a participant performed better than chance at a behavioral task. As usual, in each bootstrap step the observed data (e.g. a series of 1s and 0s for heads or tails) are resampled to obtain the alternative distribution θ_1_. However, for estimating θ_0_ we instead generate a *new* set of 1s and 0s of the same number as the observed trials using the chance probability (i.e. 0.5 or whatever the chance level is). Alternatively, one can also permute the raw trial data in each resampling step and recalculate the accuracy. The latter approach is advised when the design is unbalanced or if there is any suspicion that chance may not have a binomial distribution. In all other ways the procedure works as described.

A very similar approach for simulating a chance distribution based on the assumptions underlying the null hypothesis can also be used for other problems, for example comparing the performance of a group of participants against chance. In this case, we can simulate θ_0_ by generating a new set of ‘chance’ participants at each resampling step under the same conditions as the actual experiment (same chance probability, number of trials, and number of participants).

### Bootstrapping curve fits

The bootstrapped evidence procedure also affords itself easily for curve fitting or regression analyses. The estimated effect size in this case is the coefficient of determination, R^2^ (or goodness-of-fit). Otherwise the procedure works in much the same way as for calculating correlations. Under the null hypothesis the observations for the dependent and independent variables are scrambled randomly with replacement. Under the alternative hypothesis, the pairing is kept intact but observations are resampled with replacement.

## Results

The principle underlying the BSE test is that under both the alternative (H_1_) and the null hypothesis (H_0_) the results follow a probability distribution (Fig. 2, left panels). In classical statistics, the p-value reflects the distance of the observed effect, θ (e.g. a correlation coefficient), from the center of the null distribution. The one-tailed p-value is the area under the blue curve (null distribution) to the right of the red diamond, which denotes the observed effect.

While the null distribution depends on the sample variance, it nonetheless fails to take the *variability of the effect under H_1_* into account. The BSE estimates how distinct these two distributions are from one another by bootstrapping both H_0_ and H_1_ and quantifying Δ, the distribution of differences (Fig. 2, right panels), between them (see Methods). If the distribution is narrow and shifted away from zero this constitutes evidence for H_1_ (Fig. 2A). However, when the distribution is narrow but centered on zero this is instead evidence for H_0_ (Fig. 2B). A wide Δ distribution provides only inconclusive evidence (Fig. 2C).

The BSE is expressed by ε, which is effectively a *signal-to-noise ratio on a logarithmic scale*. It is the ratio of the observed effect, μ, and the uncertainty, σ, with which it can be estimated (Fig. 2 and Methods). When μ is smaller than σ and thus the Δ distribution overlaps zero (as quantified by ω), ε decreases as sample size, n, increases. Since ε is the logarithm of this ratio, it is positive when the data support H_1_ and negative when they favor H_0_. If ε is near zero (-0.5<ε<0.5) the evidence is inconclusive.

For the near perfect correlation with n=5 (Fig. 1A) the evidence is only ε=0.7. In contrast, for the modest correlation with n=40 the evidence ε=1 (Fig. 1B). The data in Fig. 1C,D are uncorrelated and neither correlation would reach classical significance. However, for a small sample size of n=5 (Fig. 1C) the evidence is inconclusive (ε=-0.4) while for a large sample (Fig. 1D) the evidence compellingly favors the null hypothesis (ε=-1.6).

The assumption made by parametric tests of normally distributed errors is often violated as in Fig. 1E. Even though the classical p-value is highly significant (r=-0.45, p=0.0034), the evidence for H_1_ is fairly weak (ε=0.6). The reason is that the data are heteroscedastic and thus skew classical Pearson’s correlation: the residuals of a linear fit depend on x. While y is chosen from a random normal distribution each point is also multiplied by the absolute magnitude of its paired value in x [15]. Such situations can readily occur in real experimental data: for instance, the proliferation of cell growth or the mean firing rate of neurons may also be accompanied by greater variability in those measures. This in turn could skew any correlations between these measures and an independent variable.

Notably, even increasing the sample size does not alleviate this problem. The data in Fig. 1F were drawn from the same heteroscedastic population but the sample size is five times larger (n=200) and the correlation is again highly significant (r=0.22, p=0.0014). Even robust significance tests, including those specifically developed to control for heteroscedasticity, do not fare any better (skipped correlation [22–24]: t=4.03, t_critical_=2.35; permutation test: r=0.22, p=0.0013; Spearman’s rho: ρ=0.18, p=0.0105; Kendall’s tau τ=0.14, p=0.0032; percentage bend correlation [22]: r=0.2, p=0.0042; Shepherd’s pi [25]: π=-0.23, p=0.0034). In contrast, the BSE test suggests only inconclusive evidence for H_1_ (ε=0.4) because bootstrapped distributions for H_1_ and H_0_ overlap substantially. In comparison, a homoscedastic data set with the same effect and sample size would produce more convincing evidence for H_1_ (ε=0.7).

### Performance on simulated data

For a more objective evaluation of the method I ran a series of simulations. For several sample sizes (n=15, 30, 60, 120, 240, and 480) I generated 5,000 data sets each drawn from two different distributions in which H_0_ is true: an uncorrelated bivariate Gaussian and the same heteroscedastic distribution underlying Figures 1E,F. For each simulated data set I calculated the bootstrapped evidence, ε, the classical parametric p-value and a default Bayes factor for H_1_ over H_0_ (BF_10_) [26].

Fig. 3A shows the distribution of these inferential statistics for uncorrelated Gaussian data with the various sample sizes denoted by different colors. As sample size increases the distributions for the default Bayes factor and bootstrapped evidence become increasingly shifted towards negative numbers, indicating increasing support for H_0_. In contrast, the distributions for classical p-values remain the same irrespective of sample size because under H_0_ the distribution of p-values is uniform (because the x-axis is logarithmic this manifests as a long leftwards tail). This ensures that, provided the assumptions of the test are met, the false positive rate in classical statistics is constant across sample sizes when H_0_ is true. This illustrates why classical *p-values can never provide evidence for the null hypothesis* [20]. When H_0_ is true, a proportion of tests given by the a level will be false positives. Because the estimated effect size with large sample sizes is typically very small (i.e. close to the truth of zero effect), trivially tiny effects may thus become statistically significant.

**Fig. 3.**
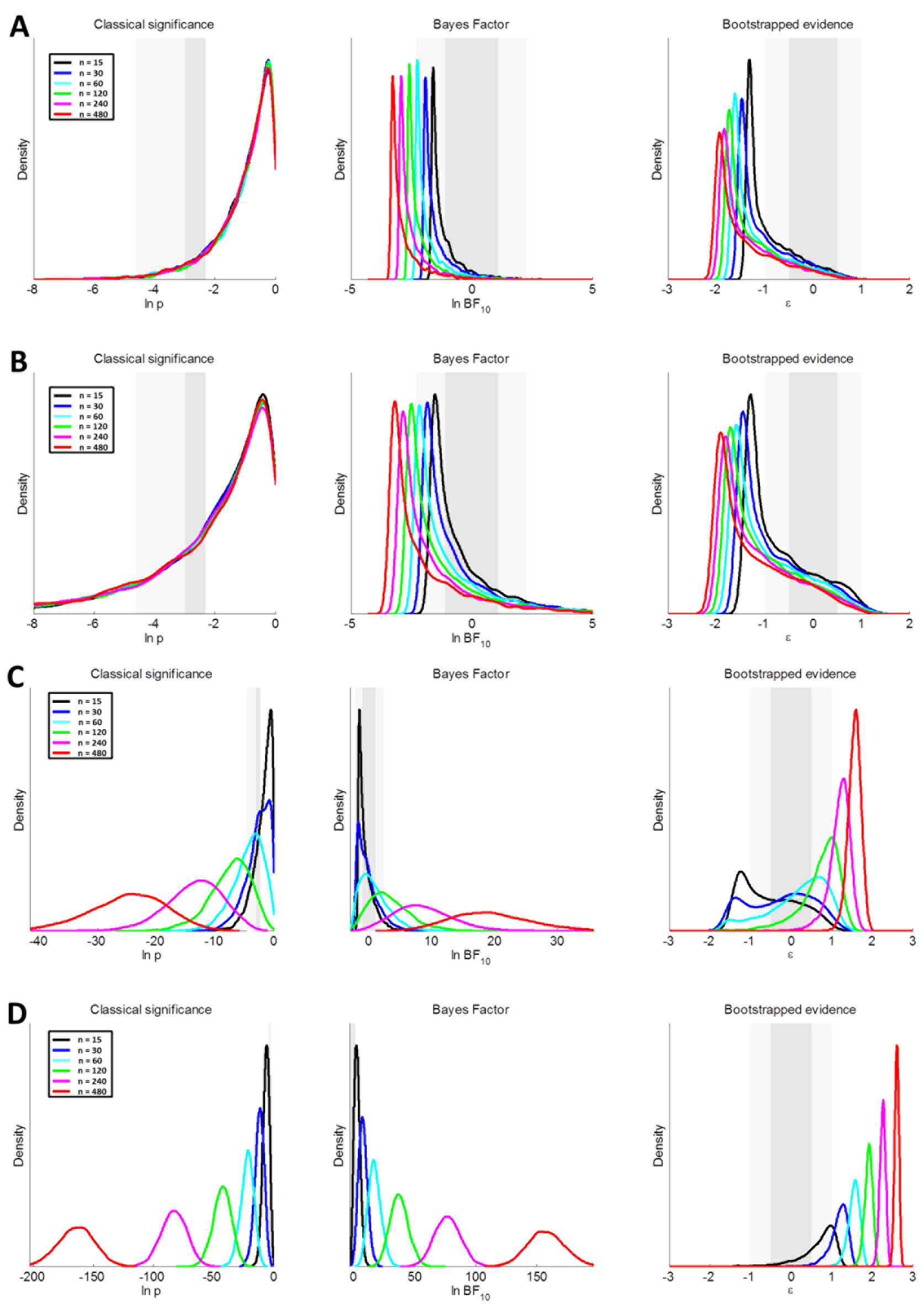
Distributions of statistical evidence in 5,000 simulations of uncorrelated Gaussian data (A), severely heteroscedastic data (B), weakly correlated data with ρ=0.3 (C) or strongly correlated data with ρ=0.7 (D). Six sample sizes were tested (see color code). Each panel shows distributions for the classical p-value (left), default Bayes factor [26] BF_10_ (middle), and bootstrapped evidence ε (right). The dark shaded regions denote “inconclusive” results (see text). The light shaded regions denote results that pass a basic criterion but provide no strong evidence for a given hypothesis.

The grey shaded regions in each panel indicate the boundaries of commonly used criterion levels. For classical statistics the dark grey region corresponds to p-values between 0.05-0.1, sometimes called “marginally significant.” The light grey region denotes the range between 0.01-0.05. Any p-value to the left of the light grey region would constitute a significant result. For Bayes factors and the bootstrapped evidence the regions are symmetric around 0. The dark grey region corresponds to inconclusive evidence that supports neither H_1_ nor H_0_ (i.e. **⅓**-3 for BF_10_, -0.5-0.5 for ε). The light grey regions refer to evidence that passed the criterion but which is still relatively weak (i.e. BF_10_ between 10^-1^ and **⅓** or 3 and 10; ε between -1 to -0.5 or 0.5 to 1). The proportion of these statistics to the right of the criterion becomes smaller as sample size increases.

Fig. 3B shows results from simulations using the uncorrelated but severely heteroscedastic distribution. For all sample sizes the distributions for classical p-values are biased so that false positives are drastically inflated. The same is also somewhat true for the default Bayes factor and bootstrapped evidence. However, for the BSE the skew is at worst modest, while the proportion of Bayes factors exceeding “strong” evidence for H_1_ is larger. This is because the default Bayes factor is a function of the Pearson’s correlation coefficient and the sample size. It is therefore skewed by heteroscedasticity in the same way as classical statistics. However, since the BSE is based on non-parametric resampling it is less affected by violations of parametric assumptions.

Next I performed a sensitivity analysis determining how well the BSE test detects true effects. I repeated the same kind of simulation but now data were chosen from a Gaussian bivariate distribution with population correlations of ρ=0.3 or ρ=0.7. Unsurprisingly, for all of the three procedures the evidence for H_1_ becomes stronger as the sample size increases. For ρ=0.3 the evidence passes criterion only for larger samples sizes (Fig. 3C) while for most data both Bayesian and bootstrapped evidence remains inconclusive. Classical statistics are less conservative as the peak of the distribution with n=120 (green curve) is already below the p<0.01 threshold. For ρ=0.7 the evidence with most sample sizes passes criterion (Fig. 3D).

I summarized the false positive and correct detection rates as a function of sample size. As expected, for classical statistics the false positive rate remains constant near the nominal level of 5% across all sample sizes, if data are Gaussian and homoscedastic (Fig. 4A). For BF_10_ and ε, false positive rates for standard criteria (BF_10_>3 and ε>0.5, respectively) decrease as sample size increases. For either method the false positive rate is already below 5% even at the smallest sample size (n=15). When heteroscedasticity is present, false positives are dramatically inflated: approximately one in four tests are positive at p<0.05 (Fig. 4B). Both evidence-based methods also show some inflation; however, false positives for the BSE are only about half that for the default Bayes factor. For the smallest sample size (n=15) the worst false positive rate for ε>0.5 is ~8.5% compared to ~14.7% for BF_10_>3. When there is a real effect (ρ=0.3) the detection rate rises steeply and then saturates for all three methods but Bayes factors and BSE are more conservative than classical p-values (Fig. 4C).

**Fig. 4.**
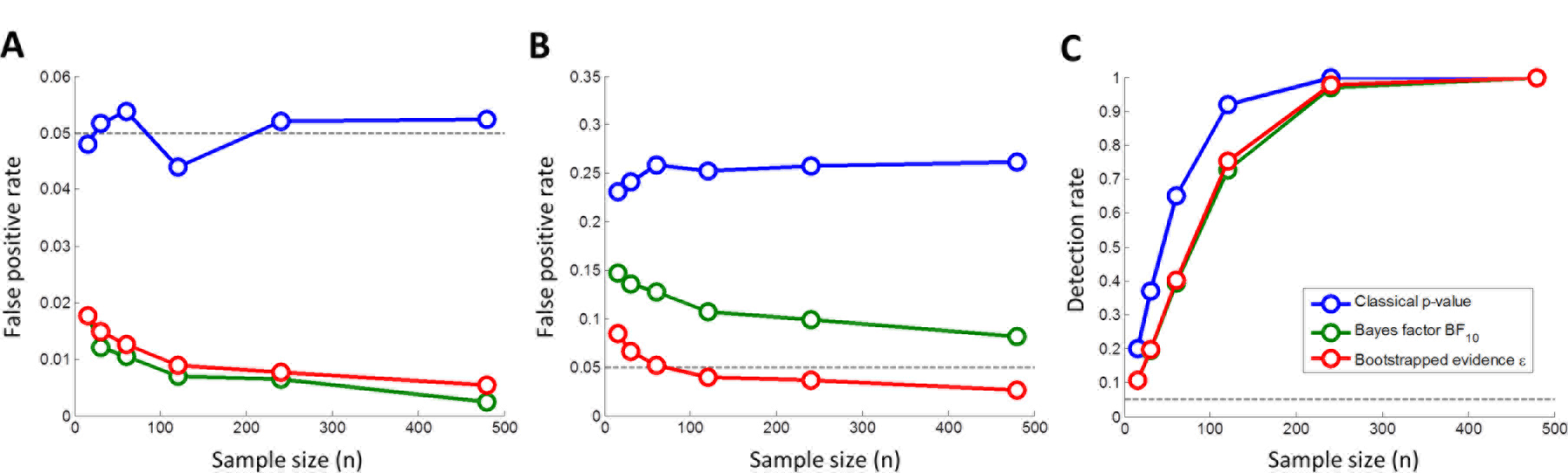
Detection rates from the simulations in Fig. 3 plotted against sample size for classical p<0.05 (blue), default Bayes factor [26] BF_10_ (green), and the bootstrapped evidence ε (red). A. Uncorrelated Gaussian data. B. Severely heteroscedastic data. C. Correlated data with ρ=0.3.

While the conclusions one would draw from all three approaches are usually similar, one notable difference is evident between classical and Bayesian inference and the bootstrapped evidence: distributions for ε tend to become *narrower as sample/effect size increase*. In contrast, the distributions for p-values and BF_10_ become wider. Note that all of these plots are on logarithmic scales (log-transformation is inherent to the calculation of ε; see Fig. 2 and Methods). Despite this, the distributions for p-values and Bayes factors display extraordinary variability, e.g. the distribution for ρ=0.3 at the largest sample size of n=480 (Fig. 3C, red curves). Here 95% of simulated p-values are between 8.5×10^-18^ and 1.8×10^-06^. All are highly significant at p<0.001 but this range spans many orders of magnitude. The default Bayes factor behaves similarly. The equivalent range spans BF_10_ between 3,212 and 378 quadrillion. Any of these would constitute “decisive” evidence of BF_10_>100 [26,27]. But from a pragmatic view, how much more confident should we be of the highest Bayes factor in this range compared to the lowest? In comparison, the equivalent range for the bootstrapped evidence is between 1.2 and 1.9. Again, these are well above even a strict criterion of ε>1 but there is no stark discrepancy between the weakest and strongest evidence. This is because rather than determining probabilities it reflects the precision with which the population effect size can be estimated. The precision increases with sample size. Thus, replicate experiments will produce very consistent bootstrapped evidence for H_1_.

Naturally, Bayesian analysis depends on the choice of a prior but typically with a range of default priors the outcome usually does not vary qualitatively [3]. Nonetheless, choosing a prior could theoretically also lead to substantial analytic flexibility, thus inflating the “researcher degrees of freedom”[28]. The BSE test on the other hand makes no assumptions beyond the resampling strategy needed for either hypothesis.

### Evidence as a function of sample size

Both classical statistics and the default Bayes factor also place undue confidence on strong effects when sample sizes are small as in Fig. 1A. The Bayes factor is rather large BF_10_=90.2 while the BSE is modest (ε=0.7). The Bayes factor reflects how much more probable the data are under H_1_ than H_0_ [29]. However, from a pragmatic perspective this could nonetheless be problematic Combining publication bias towards positive findings with underpowered experiments, high Bayes factors may thus be misinterpreted as strong evidence for the alternative hypothesis. Given the problems with spurious results and reproducibility in the scientific literature [30] this could be problematic.

Fig. 5 plots the evidence for a range of effect sizes against sample size. In most situations, the conclusions we would draw from bootstrapped evidence (Fig. 5A) and the default Bayes factor (Fig. 5B,C) are largely the same. For strong effects, the evidence rises continuously beyond the inconclusive region, while for weaker effects the evidence starts off as indistinguishable from the situation when the null hypothesis is true (black curves) until it departs and also rises. This behavior is natural because if the true effect is weaker than what could be meaningfully detected given the data at hand this constitutes support for the null hypothesis.

**Fig. 5.**
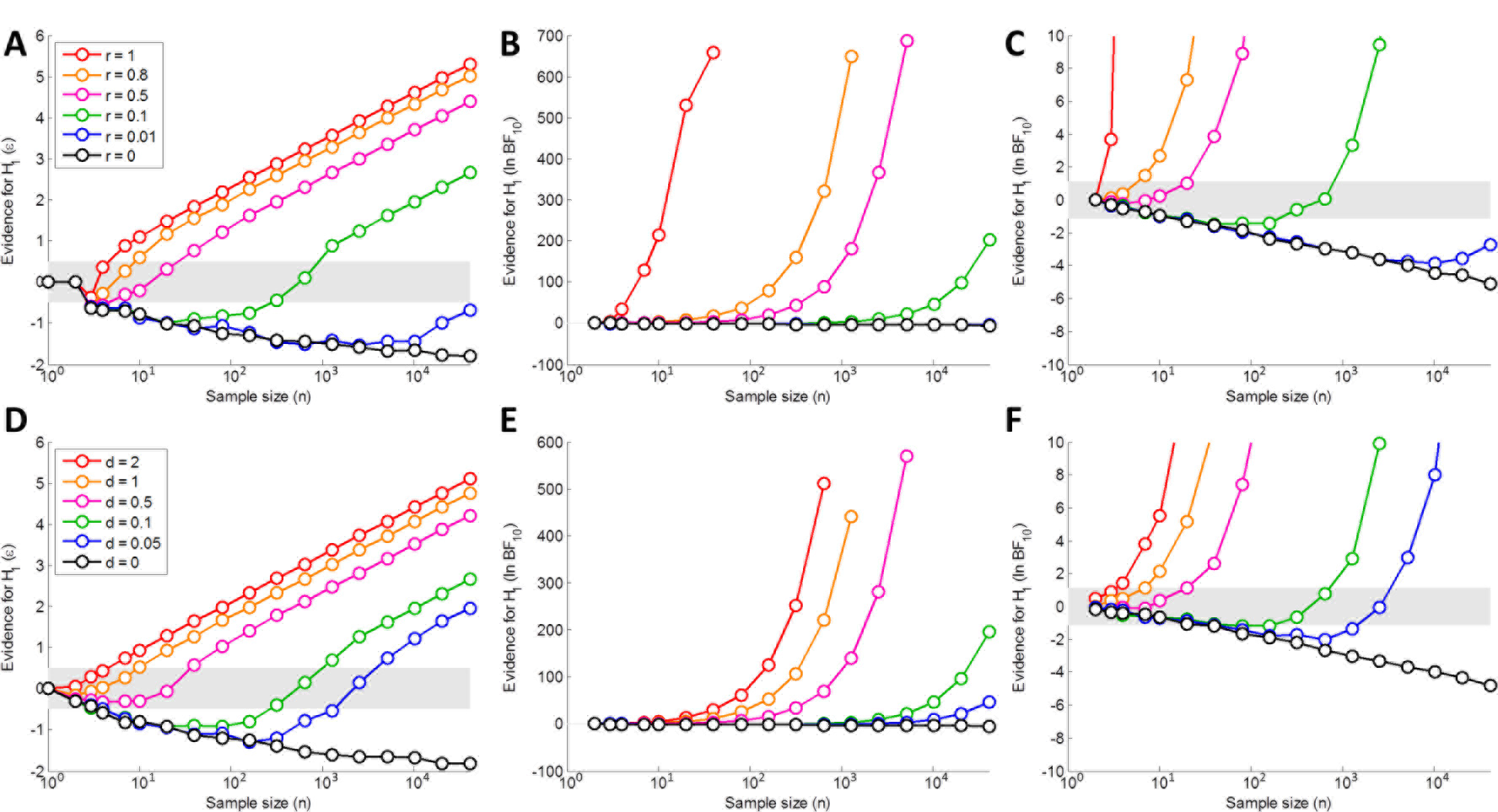
Statistical evidence for H_1_ plotted against sample size for a range of effect sizes (see color code). A-C. Correlation analysis. D-F. Comparing the means of two samples. Individual panels show the bootstrapped evidence ε (A,D) or the default Bayes factor [26] BF_10_ (B-C, E-F). Panels C and F plot the Bayes factor with y-axis zoomed in on zero. The shaded grey region denotes “inconclusive” evidence (i.e. -0.5<ε<0.5 or ⅓<BF_10_<3, respectively). For the bootstrapped evidence (A,D) these data represent the mean across 100 simulations.

The slopes of the curves for the bootstrapped evidence are far less steep. Thus it is possible to see the behavior for the full range of conditions within the same plot. There is however one considerable difference: for a perfect correlation (ρ=1) the default Bayes factor immediately rises even at tiny sample sizes (Fig. 5C). At n=3 the BF_10_ is already 48.8. In contrast, the BSE for this point is low (ε=-0.4) and inconclusive. As sample size increases, so does the bootstrapped evidence. At n=4 it is still inconclusive but favoring H_1_ (ε=0.4). At n=7 it clearly supports H_1_ (ε=0.9) and it continues to rise as sample size increases. This behavior is more intuitive than that of the default Bayes factor and also the classical p-value, which would be extremely significant in all these situations. Compare this to the earlier example of a strong correlation (r=0.9953) with a small sample size of n=5 (Fig. 1A). Classical inference would be extremely significant (p<0.001) and the default Bayes factor would yield “very strong” evidence for H_1_ (BF_10_=90.2). The BSE is however only fairly modest, especially given the strong effect (ε=0.7). It is above the criterion for conclusive evidence but it does not instill undue confidence in H_1_.

The data in this example were in fact drawn from an uncorrelated Gaussian distribution so the null hypothesis was true. The bootstrapped evidence provides an intuitive measure of the weakness of the evidence in such situations and should thus be a safeguard against weak or inconclusive results.

### Simulations of optional stopping

The BSE test has further advantages over classical inference based on significance thresholds. In classical statistics, even when there is no true effect, it is theoretically possible to reach an arbitrarily significant p-value, if data collection continues until the p-value passes the significance threshold. This is known as “optional stopping”, which is an incorrect but possibly widespread use of classical statistics [4,31]. Under the classical framework one should first define the expected effect size *a priori*, perform a power analysis to see how large a sample is needed to detect this effect with sufficiently high probability, and then collect those data without stopping until the sample is complete. However, typically this is not realistic as one can often only make a vague guess about the expected effect size.

The bootstrapped evidence does not suffer from this conundrum. First, even if a dubious optional stopping strategy is used, the false positive rate is not inflated substantially. I simulated this 1,000 times by drawing data repeatedly from an uncorrelated bivariate Gaussian distribution thus successively increasing the sample size by 1, starting with a minimal sample of n=5. At each step I applied classical statistics, the default Bayes hypothesis test [26], and the BSE test. The first instance one of these tests passed the criterion level, that is p<0.05 for classical statistics, BF_10_>3 or BF_10_<⅓ for Bayes factors, and ε>0.5 or ε<-0.5 for the bootstrapped evidence, I recorded the measure of evidence. In addition, I also performed this procedure on the bootstrapped uncertainty and recorded the first instance that σ<0.2. If none of the measures reached criterion simulated data collection would cease at n=150.

Under the assumptions of classical statistics this false positive rate should be near 5%, however, the actual probability was much greater, 36.4%, illustrating the considerable problems optional stopping can cause in classical inference. In contrast, the false positive rates of the evidence-based methods was much lower and well below the classical a level (Bayes factor: 2.9%; bootstrapped evidence: 2.8%).

I repeated the same simulation but this time drawing from the heteroscedastic distribution used in previous examples (Figures 1E,F). Now classical statistics massively inflated support for the alternative hypothesis with a false positive rate of 84.6%. Default Bayes factors fared a lot better but are nonetheless strongly skewed (20.6%). For the bootstrapped evidence on the other hand the false positive rates were only half that (10.3%), again reflecting the fact that it is based on a non-parametric procedure that takes into account the anomalous distribution of the data.

This demonstrates that optional stopping based on symmetric evidence is far less problematic than for classical statistics. In particular, sequential analysis until the bootstrapped evidence reaches conclusive support for either H_1_ or H_0_ results in only minimal false positive rates even in extreme situations. However, there is an even better optional stopping strategy that could be employed in the bootstrapped evidence framework. When data collection continued until the bootstrapped uncertainty, σ, was 0.2, the false positive rate for using ε>0.5 in the first scenario (homoscedastic Gaussian data) was only 1.3%, while for the heteroscedastic data it was 7.9%. This suggests that using a criterion uncertainty level is the most optimal strategy for minimizing spurious findings in sequential analysis.

### Example 1: Anscombe’s quartet

Simulations are crucial for testing a method’s performance because the ground truth is known. However, for illustration I also apply the method to Anscombe’s quartet [32] a famous demonstration of the pitfalls of correlation analysis. It consists of four data sets, each comprising 11 pairs of variables, in which Pearson’s correlation produces (almost) identical results (r=0.82, p=0.002). Applying the BSE test reveals that while the data afford low but sufficient confidence for the correlation in the first three data sets (Fig. 6A-C), in the final example (Fig. 6D) the evidence clearly supports H_0_ (ε=-1.4) because one influential outlier drives the correlation but the remaining data are uncorrelated.

**Fig. 6.**
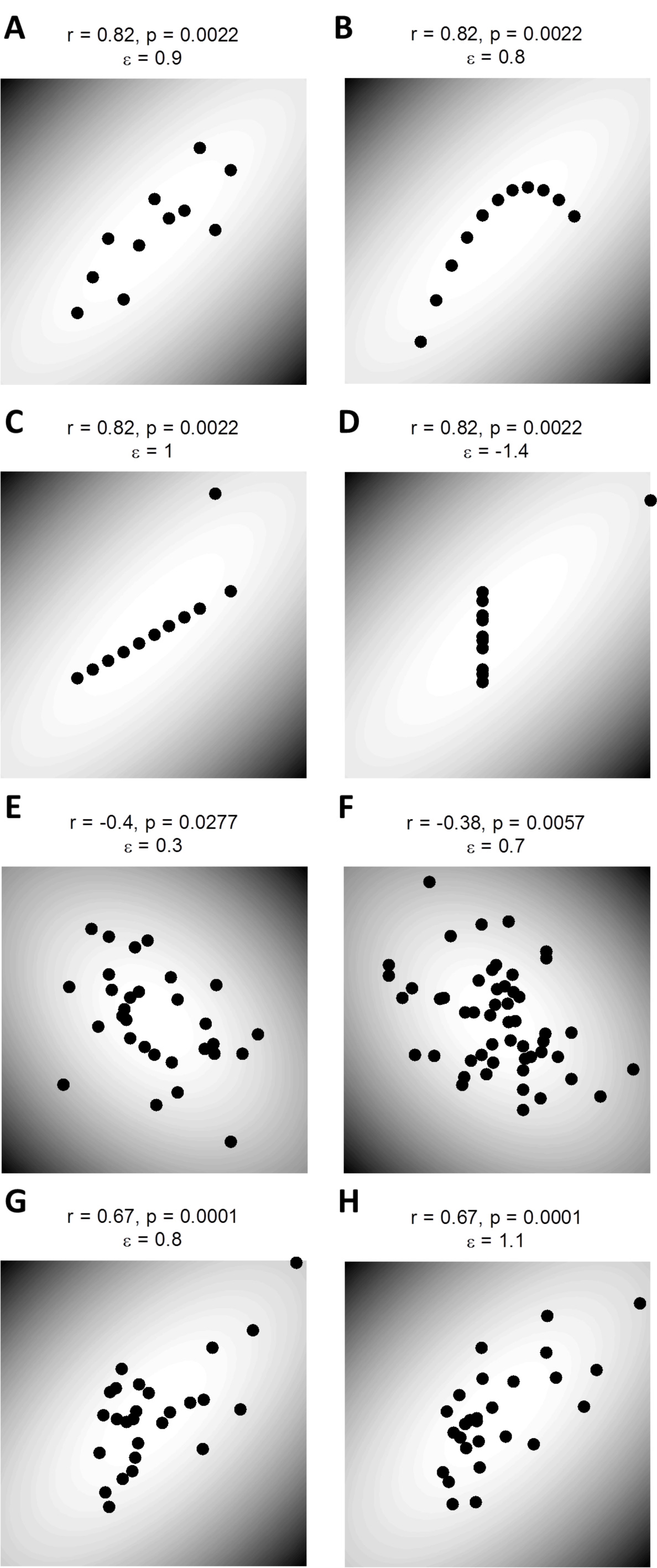
Example data sets. A-D. Anscombe’s quartet: Each data sets has approximately the same Pearson’s correlation (r=0.82, p=0.0022) and thus all have a default Bayes factor [26] BF_10_≈23. Panels show Typical Gaussian data (A), data showing a perfect non-linear relationship (B), a perfect correlation contaminated by one influential outlier (C) and uncorrelated data contaminated by one influential outlier (D). E-H. Experimental data showing correlations between visual cortical surface area and perceptual function. Correlations between V1 area and Ebbinghaus illusion strength from [33] (E) and [34] (F). Correlations from [35] between travelling wave speed in binocular rivalry and the surface areas of V1 (G) and V2 (H). All other conventions as in Fig. 1.

Interestingly, the confidence in H_1_ is actually subtly greater (ε=1) for the third example (Fig. 6C) than the first (ε=0.9, Fig. 6A). This is because a single outlier contaminates the perfect correlation in this example, whereas the first example contains noisy but normally distributed data.

The BSE does not distinguish strongly between the first and second examples (Fig. 6A,B). The second example contains a perfect relationship between x and y; however, it does not conform to the linear relationship assumed by Pearson’s correlation. Curve fitting can also be implemented in the BSE framework (see Methods). Here we could compare a simple linear fit to polynomial curves. The evidence for H_1_ with a second-order polynomial is considerably greater (ε=1.6) than for a standard linear model (ε=1). Interestingly, the BSE is also robust to overfitting more complex models: the evidence for higher-order polynomials is weaker than for the second-order (third-order: ε=1.3; fourth-order: ε=1).

### Example 2: Links between visual cortex surface area and perceptual function

I further applied the BSE test to published experimental data that showed correlations between the size of early visual areas and perceptual function. These studies hypothesized that the transmission speed/strength of lateral connections running tangential to the cortical surface is reduced for individuals with larger cortical surface areas. In the first two studies, this should manifest as an anti-correlation between the strength of the Ebbinghaus illusion and V1 surface area [33,34]. Classical statistics confirmed this hypothesis in both studies (Fig. 6E,F). However, according to the BSE the findings of the initial study were inconclusive (r=-0.4, p=0.028, ε=0.3). The second study used a more sophisticated design producing more compelling evidence for this link (r=-0.38, p=0.006, ε=0.7; note, however, that this study also normalized V1 area by the whole cortical surface area to control for non-linearity issues and other confounds. For the sake of consistency with the other findings I chose not to apply this correction here).

The third study [35] reported a linear relationship between the speed of travelling waves in binocular rivalry and the surface areas of V1 and V2. Classical statistics were very similar for both regions (r=0.67, p=0.0001). However, the bootstrapped evidence was in fact lower for V1 (Fig. 6G; ε=0.8) than V2 (Fig. 6H; ε=1.1), possibly because influential outliers affected the former.

### Other statistical tests

The BSE test can also address many other questions. One simply needs to change how the effect size is calculated and how data are resampled during bootstrapping (see Methods). For example I also ran simulations for comparing the means of two samples (Fig. 5 and Supplementary Fig. 1).

**Supplementary Figure 1.**
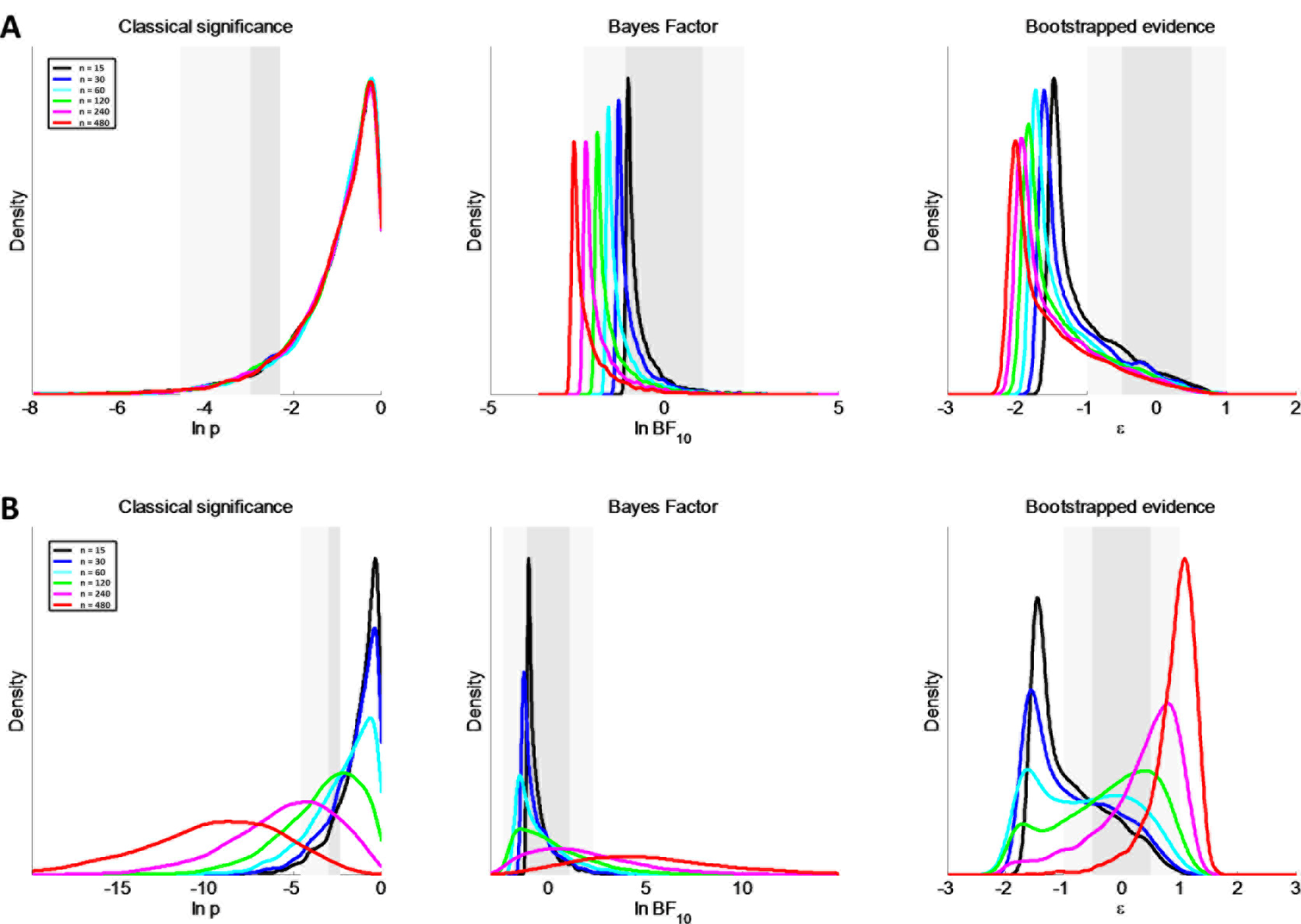
Distributions as shown in Figure 4 but for comparing the means of two samples. A. No difference, i.e. normally distributed data with unit standard deviation and a population mean of 0. B. A weak effect with a population difference of 0.25 and unit standard deviation. All other conventions as in Figure 4.

### Example 3: reassessing evidence for precognition

A few years ago a psychology study reported experimental evidence for the proposition that participants had precognitive abilities [16]. This study was criticized as an illustration of the shortcomings of classical inference: Bayesian reanalysis found little evidence in favor of precognition [3]. However, a Bayesian analysis by the original author supported his claims [17]. The conclusions are evidently dependent on the exact prior chosen [3,18]. A cautious prior seems advisable when evaluating such unusual claims but the debate illustrates how Bayesian inference can appear to lack objectivity.

Here I reevaluate claims from a more recent study on precognitive abilities [36] with the BSE test (see Methods). It is a perfect test of the method because these results seem biologically and physically implausible but both classical and default Bayesian inference [3] nonetheless support the alternative hypothesis for several of the studies (Table 1). In contrast, the BSE does not provide convincing support for precognition. It is noteworthy that despite a large sample size of 1222 participants, even the web-based study 3 only produced inconclusive evidence (ε=-0.1). This illustrates that in comparison to classical inference, the BSE is far less susceptible to the inflation of “significant” findings with large sample sizes. Taken together this suggests that the available data do not provide conclusive evidence for either H_1_ or H_0_.

**Table 1.** Reanalysis of data purportedly showing precognitive abilities [36] and a simulated example of a clinical trial. For each experiment this shows the result using classical statistics, a default Bayes factor [20] and the bootstrapped evidence ε. For Bayes factors and bootstrapped evidence verbal descriptions of the strength of evidence for H_1_ and H_0_ are also given. Cells with bold font denote tests formally supporting H_1_, that is, if evidence for the alternative hypothesis is above criterion (i.e. p<0.05, BF_10_>3 or ε>0.5). All these statistics are based on my own analysis of the raw data. The asterisks indicate that the data for these studies were simulated based on the reported effect and sample sizes because the raw data were not available to me.

| Experiment | Classical statistics | Bayes Factor, $BF_{10}$ | Bootstrapped evidence, $\epsilon$ |
| --- | --- | --- | --- |
| Maier Study 1 | **t(110)=-2.65, p=0.0092** | 2.9 (anecdotal $H_1$) | 0.2 (inconclusive) |
| Maier Study 2 | **t(200)=-2.99, p=0.0032** | **5.9 (moderate $H_1$)** | 0.3 (inconclusive) |
| Maier Study 3 | **t(1221)=-2.36, p=0.0186** | 0.5 (anecdotal $H_0$) | -0.1 (inconclusive) |
| Maier Unsuccessful 1 | t(62)=0.16, p=0.8700 | 0.14 (moderate $H_0$) | -1.8 (compelling $H_0$) |
| Maier Unsuccessful 2 | t(405)=0.44, p=0.6587 | 0.06 (strong $H_0$) | -1.8 (compelling $H_0$) |
| Maier Unsuccessful 3 | t(639)=-1.3, p=0.1943 | 0.1 (strong $H_0$) | -0.3 (inconclusive) |
| Maier Study 4 | t(326)=-1.81, p=0.0717 | 0.3 (strong $H_0$) | -0.2 (inconclusive) |
| Aspirin study* | **t(21998)=5.19, p<10<sup>-6</sup>** | **10561.9 (decisive $H_1$)** | **1.3 (compelling $H_1$)** |

Is the BSE test simply too conservative to reveal these rather subtle effects? To test whether the low evidence in my reanalysis could be due to a lack in sensitivity, I also used the BSE test to evaluate a small effect size in the clinical literature. In this study, the effect of aspirin on cardiac health was tested in a large sample (n≈22,000) [37]. The effect was minute (Cohen’s d≈0.07) but highly significant, and it was thus deemed to be of practical value in promoting health. In the absence of raw data I simulated this data set using the reported effect and sample sizes. Here the BSE test agrees with classical statistics and Bayesian inference: the observed data strongly support (ε=1.3) the efficacy of the drug (Table 1). The sample size in this case is more than sufficient to conclude that this effect is *small but real*. In contrast, to provide conclusive evidence for the tiny precognition effects, the sample sizes would have to be orders of magnitude larger (Fig. 5D).

## Discussion

Here I outlined the bootstrapped evidence test, which makes minimal assumptions and is easily applicable to numerous situations. It quantifies non-dichotomously the evidence *for the alternative or the null hypothesis* rather than whether a particular statistic passes an arbitrary significance threshold. A result does not stand or fall based on its exact value. Rather it allows us to express how convincing a result is. This also allows for the use of sequential sampling strategies, which can greatly benefit research practice especially when there is uncertainty about the expected effect size.

A possible criticism of non-dichotomous alternatives to null hypothesis significance testing is that any measures could be subject to the same thresholding dilemma as classical p-values. If the consensus emerges that ε>0.5 is sufficient evidence for H_1_ then are we not merely shifting the problem from p-values to a different measure? However, even in the classical framework many researchers regularly make non-dichotomous judgments based on p-values. “Marginally significant” findings are often reported that do not quite pass reach p<0.05. Many researchers are probably more convinced by p=10^-17^ than p=0.049. The bootstrapped evidence test directly quantifies the reliability of the data in drawing conclusions about the two competing hypotheses. Being a new measure it will make it easier for researchers to adopt non-dichotomous thinking than with p-values.

It remains unclear in how far researchers need dichotomous thresholds for statistical inference and whether labels for the strength of evidence, such as those employed for Bayes factors [26,27], are necessary. I would argue that they are not and deliberately refrained from proposing a categorical stratification of ε. For replication attempts or incremental experiments for which clear predictions can be made, ε between 0.7-1 can already be very convincing. In contrast, for more extraordinary claims even ε=1 is still low.

Another problem with most commonly used statistics is that they are based on parametric assumptions that may not hold. Anomalous data can skew the default Bayes factor to a similar extent as inferences based on classical p-values, because it is based on the same effect size calculations. Robustness to outliers and heteroscedasticity could be incorporated into Bayesian hypothesis testing, e.g. by outlier removal procedures. Within the framework of classical inference robust hypothesis testing suggest following a complicated tree of tests for the presence of outliers and heteroscedasticity, and then applying the appropriate robust test depending on the situation [22,23]. This is often accompanied by remonstrations that there is a “statistical toolbox” that should be employed and that one method is not best for every situation [9].

While it is doubtless true that inference should never be made without thought [10], it is nevertheless advisable to seek a method that serves a universal purpose because in practice many researchers are not statisticians. The idea of a “statistical toolbox” is fraught with danger. It must inevitably result in increasing the underreported flexibility in the range of methods employed by published studies [28,38].

Of course the BSE test does not preclude the use of other statistical methods. In particular, I view it as a *complement to Bayesian inference* rather than an alternative. It provides a common starting point for inference. It is an objective method with minimal assumptions that in principle works exactly the same for almost any situation. The BSE test is particularly useful when there is substantial uncertainty about the expected effect size, when there are violations of parametric assumptions, and generally whenever application of Bayesian inference is difficult. It is also especially useful for exploratory analyses. The bootstrapped distribution of the effect size estimate could inform a prior used for Bayesian analysis of subsequent replication attempts.

The BSE framework also directly encourages sharing of raw data. While it can be applied to data simulated based on parameter estimates, as I have done for the Aspirin study in Table 1, this neglects additional information about potential data anomalies (Fig. 6). For the purpose of metaanalysis or even simply reanalyzing individual research findings, the full advantages of the BSE test become apparent when it is used on raw data.

Finally, it is important to remember that no statistical procedure can replace scientific scrutiny. Statistics do not confirm or refute theories. No amount of statistical evidence can prove whether particular phenomena, be it social priming, brain-behavior correlations or even far-fetched claims like precognition, exist. They can only provide support that the predictions a particular hypothesis makes are likely not to have occurred by chance or another, more trivial explanation.

## Acknowledgements

I thank Ged Ridgway, Benjamin de Haas, and Micah Allen for comments on previous versions of this manuscript.

